# Comparative transcriptomic analysis reveals translationally relevant processes in mouse models of malaria

**DOI:** 10.1101/2021.05.11.443626

**Authors:** Athina Georgiadou, Claire Dunican, Pablo Soro-Barrio, Hyun Jae Lee, Myrsini Kaforou, Aubrey J. Cunnington

**Affiliations:** Section of Paediatric Infectious Disease, Department of Infectious Disease, Imperial College London, London, UK; Centre for Paediatrics and Child Health, Imperial College London, London, UK; Institute for Molecular Bioscience, University of Queensland, Brisbane, Queensland, Australia

## Abstract

Recent initiatives to improve translation of findings from animal models to human disease have focussed on reproducibility but quantifying the relevance of animal models remains a challenge. Here we use comparative transcriptomics of blood to evaluate the systemic host response and its concordance between humans with different clinical manifestations of malaria and five commonly used mouse models. *Plasmodium yoelii* 17XL infection of mice most closely reproduces the profile of gene expression changes seen in the major human severe malaria syndromes, accompanied by high parasite biomass, severe anemia, hyperlactatemia, and cerebral microvascular pathology. However, there is also considerable discordance of changes in gene expression between species and across all models, indicating that the relevance of biological mechanisms of interest in each model should be assessed before conducting experiments. Our data will aid selection of appropriate models for translational malaria research, and the approach is generalizable to other disease models.

## Introduction

Animal models have played an important role in current understanding and treatment of many human diseases. Historically animal models were often selected because they reproduced certain clinical or pathological features of human disease (1), and their use has often been reinforced when treatments effective in the model were found to be effective in humans. However, this approach has limitations, because the same clinical or pathological features can occur as a result of different biological processes, and mechanisms which may be important in human disease might not be recapitulated or may be redundant in animal models, and vice-versa (2, 3). A fundamental and largely unresolved question is how best to define or quantify the relevance of any given animal model to the corresponding human disease (3, 4).

Mice are the most widely used model animals for many diseases, including infectious diseases, and for study of corresponding protective or pathogenic immune responses. Mouse models have significantly broadened our understanding of the function and structure of mammalian immune systems and disease mechanisms. Despite evolutionary distance between human and mouse (5) and the high evolutionary pressure on immune systems (6), the principles of the immune systems for these species remain remarkably similar. However, there are also numerous differences between mice and humans in terms of their response to infection (5). Therefore, it is inevitable that mouse models of infection will not recapitulate all features of the human response, and this should be taken into account when using models to make inferences about mechanisms of human disease. Recently it has been proposed that unbiased approaches to assessment of the host response to infection, such as comparison of transcriptomic responses, might provide a meaningful way to quantify similarities between mouse models and human disease, to assess the relevance of the models, and to aid the selection of the best models for specific hypothesis testing (7).

The relevance of mouse models for translational research on the pathogenesis of severe malaria has been particularly controversial and has polarized the malaria research community (8). There are many different mouse malaria models, with very different characteristics dependent on the combination of parasite species (and strain) and mouse strain which are used (9). Superficially these models can, between them, reproduce almost all the clinical manifestations of human severe malaria (SM), such as coma, seizures, respiratory distress and severe anemia (10). Nevertheless, there are also notable differences to human disease, such as the lack of the pathognomonic cytoadhesive sequestration of large numbers of parasite infected red cells in the cerebral microvasculature in mice with cerebral malaria-like illness (11) (termed experimental cerebral malaria, ECM). Many host-directed treatments for SM have been effective in mice, but none have yet translated into benefit in human studies, which has been considered by some as further evidence that mouse models are of little relevance to human disease (12). We contend that this polarization of views is unhelpful, and that mouse models are likely to be useful for understanding human malaria, so long as they are used selectively with full recognition of their limitations.

In order to provide a more quantitative framework to understand how well mouse malaria models recapitulate the biological processes occurring in human malaria, and to aid selection of the most appropriate models for study of specific mechanisms of disease, we present an unbiased investigation of the similarities and differences in the host response between human malaria and mouse models using comparative transcriptomics. We demonstrate that this approach allows us to identify mouse models with the greatest similarity of host response to specific human malaria phenotypes, and that models selected in this way do indeed have similar clinical and pathological features to those of the corresponding human phenotype. We propose that this approach should be applied more broadly to the selection of the most relevant animal models for study of malaria and other human diseases.

## Results

### Mouse models of malaria

The five rodent malaria parasite species used in this study produce different kinetics of parasitemia, different rates of progression of illness (Fig. 1), and different disease manifestations. 8-week-old C57BL/6 mice infected with *P. berghei ANKA, P. yoelii 17XL* and *P. berghei NK65* developed severe illness with ascending parasitemia, consistent with previously reported outcomes of these lethal infections (12–15). Humane endpoints were reached at day 8-9 in *P. berghei ANKA*, day 5 in *P. yoelii 17XL*, and day 20 in *P. berghei NK65*. Mice infected with *P. berghei* ANKA showed typical features of ECM as assessed by Rapid Murine Coma and Behavior Scale (RMCBS) scores < 12 (13) and by histopathology. Mice infected with *P. yoelii 17XL* developed a rapidly progressive, severe infection with hyperparasitemia (14). Mice infected with *P. berghei* NK65 developed a biphasic illness with a transient recovery of initial weight loss before progression to fatal outcome in a second phase (15).

**Figure 1:**
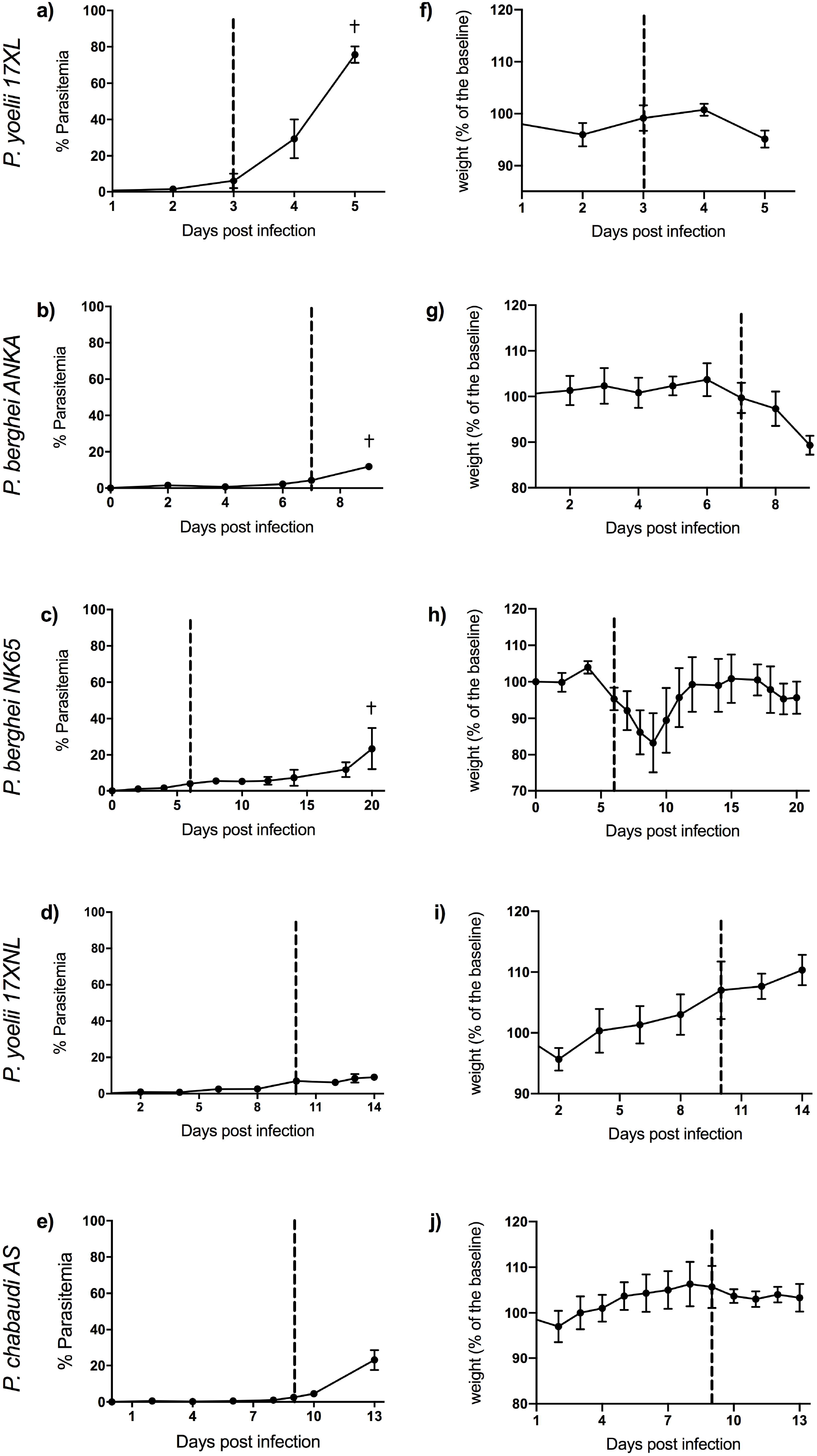
Course of infection in five mouse malaria models. Comparison of parasitemia (a, b, c, d, e) and change in weight (as percentage of baseline weight) (f, g, h, i, j) for 8-week-old C57BL/6 female wild type mice infected with: *P. yoelii 17XL, P. berghei ANKA, P. berghei NK65, P. yoelii 17XNL*, and *P. chabaudi AS, respectively*. Points show mean, and bars show SD, for n=6 mice (up to and including time point of first signs of ill health, dashed vertical line) and n=3 mice (after dashed vertical line) for each infection. † indicates humane endpoint for lethal infections.

8-week-old C57BL/6 mice infected with *P. yoelii 17XNL* and *P. chabaudi AS*, which lead to self-resolving infections, developed only mild symptoms as expected (16),(17). Maximum severity was reached around day 14 in *P. yoelii* 17XNL and day 13 in *P. chabaudi AS*.

### Comparative analysis of infection-associated changes gene expression

To objectively assess how similar disease-associated systemic processes occurring in mouse malaria models are to those occurring in human *P. falciparum* malaria, we used a comparative transcriptomic approach focussed on blood. Rather than directly comparing expression of orthologous genes in humans and mice, which would be confounded by species-specific differences in constitutive gene expression, we first identified differentially expressed genes in pairwise within-species comparisons and then used these differentially expressed genes as the basis for between-species comparisons (Fig. 2a). This also enabled us to conduct within-species adjustment for variation in leukocyte cell mixture, which is an important confounder in whole blood gene expression analysis (18). Additionally, this allows for the removal of platform-specific effects, which is especially relevant for comparisons between Microarray and RNA-Seq generated differentially expressed genes.

**Figure 2.**
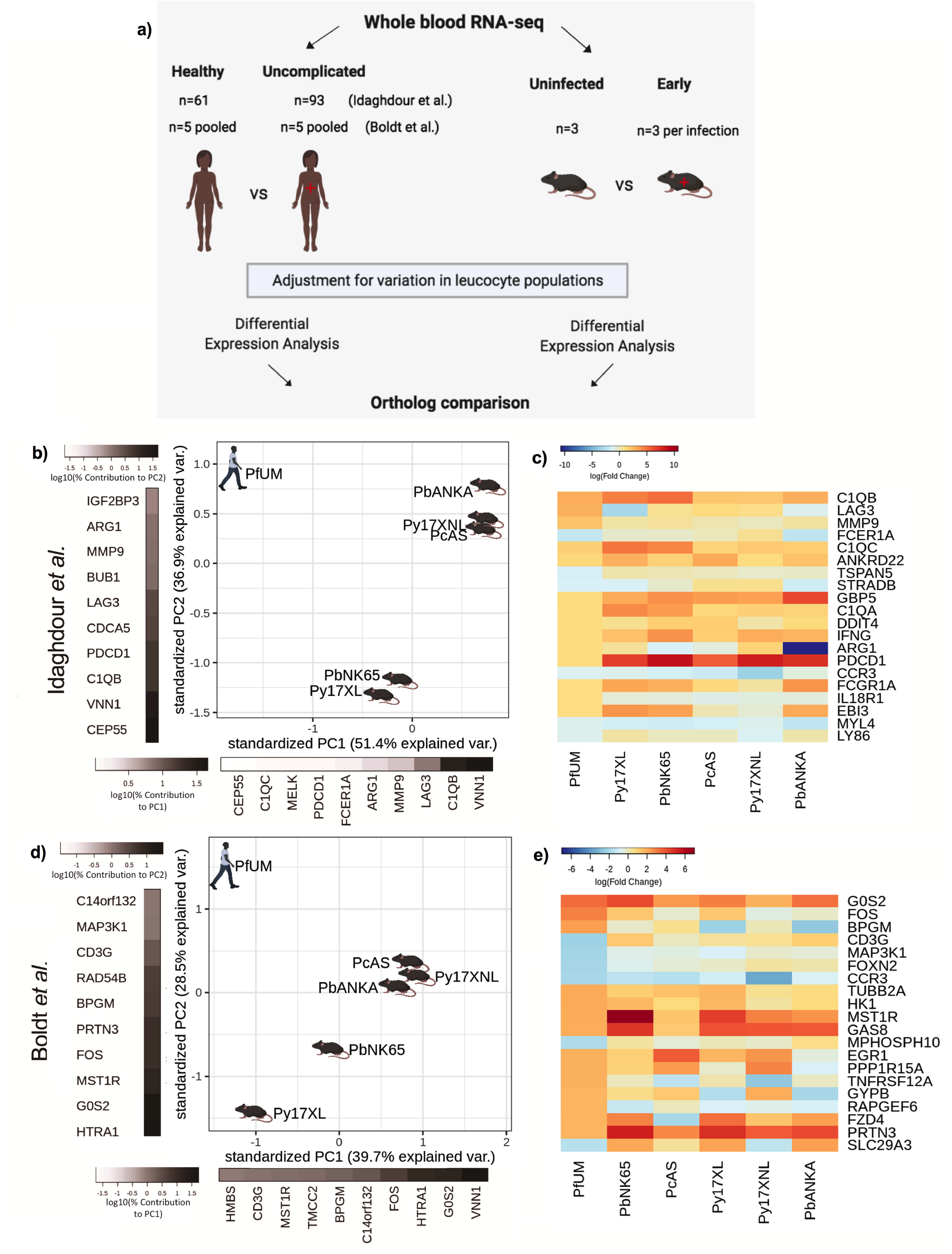
Comparison of host differential gene expression in human uncomplicated malaria and early-stage illness in five mouse malaria models. a) Schematic illustration of the comparative transcriptomic analysis. b, d) Principal component analysis (PCA) plots generated using rank-normalised log-fold change values from the human and mouse differential expression analyses. Only genes with 1:1 mouse and human orthologs and with absolute logFC value greater than 1 in the corresponding human comparison were included. Comparison of changes in gene expression in the mouse models (uninfected vs early in infection) with those in uncomplicated malaria vs healthy (PfUMH) Beninese children (b, Idaghdour et al. (19)) or Gabonese children (d, Boldt et al. (20)). The percentage of the total variation explained by principal components 1 and 2 are shown in the axis labels. Greyscale heatmaps parallel to each axis show the contributions of the 10 genes contributing most to the corresponding PC. c, e) Heatmaps show logFC for the 20 genes with the greatest absolute log fold change values in the human differential gene expression analysis, and their orthologs in each mouse model, corresponding to the analyses illustrated in b and d, respectively. Mouse models are ordered left to right in order of increasing dissimilarity to the human disease, based on the Euclidian distance calculated from all principal components (Supplementary File 8). The rows (genes) are ordered by absolute log fold change in the human comparison in descending order. n=3 for early and n=3 for late time point in each mouse model; n=93 UM, n=61 controls (Beninese children, Idaghdour et al.), n=5 pools UM and n=5 pools healthy control samples (each pool contained RNA from 4 Gabonese children with the same phenotype, Boldt et al.). The mouse model abbreviations are as follows: PbNK65 (*P. berghei NK65*), PbANKA (*P. berghei ANKA*), PcAS (*P. chabaudi AS*), Py17XL (*P. yoelii 17XL*) and Py17XNL (*P. yoelii 17XNL*).

Sometimes we may wish to investigate the host immune response to infection *per se* or alternatively we may want to investigate the processes associated with severe disease pathogenesis, and these different aims require different comparator groups. In the former situation, changes in gene expression associated with infection *per se* are best characterised by comparison between healthy and infected states, whereas in the latter situation it may be more appropriate to compare severe and non-severe infection states.

To investigate concordance of the host response to uncomplicated malaria (UM) in humans and mice, we first focussed on comparisons between subjects with UM and healthy uninfected subjects. To assess changes in gene expression due to *P. falciparum* malaria we used two human transcriptomic datasets previously published by Idaghdour et al. (19) and Boldt et al. (20), each of which included a healthy uninfected group and an uncomplicated *P. falciparum* malaria group (Supplementary File 1). For the mice, we identified changes in gene expression occurring between mice culled at first onset of clinical signs of illness and healthy uninfected control mice. To remove the confounding effect of constitutive and infection-induced differences in leukocyte population proportions in blood, all differential expression analyses were performed with adjustment for the proportions of the major leukocyte populations in blood (see Methods; Supplementary File 2). Genes with absolute log-fold change in expression >1 in the human healthy control vs UM comparison (Supplementary Files 3 and 4) and mouse orthologs (Supplementary File 5) were used for comparison between species.

First, considering only whether genes were up-regulated or down-regulated by infection in the mouse models, we found the mouse models varied from 58 to 73% concordance (Supplementary File 6, Fig. 2) with the up- or down-regulation in the human subjects in the study by Idaghdour et al. However, we reasoned that the relative magnitude of changes in gene expression is also important to identify the mouse models which most closely recapitulate the changes in gene expression in human malaria. To assess this, we weighted genes to account for the relative magnitude of change in expression in human malaria, and then performed a principal component analysis with these weighted changes in expression (see Methods). This revealed variation between the mouse models, but no model was clearly much more representative of the changes in gene expression in human UM than any other (Fig. 2b). Indeed, when we focused only on the expression of the twenty most differentially expressed genes in human UM, we found that the mouse models showed broadly similar patterns of changes in gene expression (Fig. 2c). When we examined the concordance of up- and down-regulation of gene expression between the mouse models and human malaria in the Boldt et al. dataset, we found less overall similarity between species in the direction of changes in gene expression (Supplementary File 6). Despite this, and different genes driving the axes of variation, the PCA plots revealed a remarkably similar pattern to the analysis based on the Idaghdour et al. dataset, and none of the mouse models appeared to be clearly more representative of human UM than any other when accounting for the magnitude of changes in expression (Fig. 2d). Considering the most differentially expressed genes, there was more heterogeneity in the pattern of expression (Fig. 2e) which may be partly explained by the substantially smaller size, and analysis of pooled samples in the Boldt et al. study.

Gene ontology (GO) analysis was used to examine the genes driving the axes of variation between humans and mouse models in the PCA plot. For the Idaghdour et al. dataset we found that PC1 showed enrichment of leukocyte mediated immunity and adaptive immune response, while PC2 showed enrichment for intrinsic apoptotic signalling in response to oxidative stress and regulation of T cell activation (Supplementary File 7). For the Boldt et al. dataset comparison, we found that that PC1 showed enrichment of cytokine-mediated signaling pathways and hemopoiesis as the top GO terms, while for PC2 the top GO terms included immune system process and myeloid cell development (Supplementary File 7).

### Comparative analysis of severe malaria-associated changes in gene expression

A common approach to identify processes associated with the pathogenesis of severe infection is to compare individuals with severe manifestations against other individuals who have the same infection but have not developed severe illness (7). This approach is expected to enrich for genes involved in the pathogenesis of severe illness from amongst the larger set of genes involved in the overall response to infection (7). Therefore, we identified changes in gene expression in mice between the first time point at which mice developed signs of illness (early) and the maximum severity (late time point) of each of the 5 infection models. We compared these changes in gene expression in mice with those we had previously identified in Gambian children with UM and three different severe *P. falciparum* malaria (SM) phenotypes (hyperlactatemia (HL), cerebral malaria (CM), or the combined phenotype of hyperlactatemia with cerebral malaria (CH)) (18) (Fig. 3a). To remove the confounding effect of constitutive and infection-induced differences in leukocyte population proportions in blood, all differential expression analyses were performed with adjustment for the proportions of the major leukocyte populations in blood (see Methods; Supplementary File 9).

**Figure 3:**
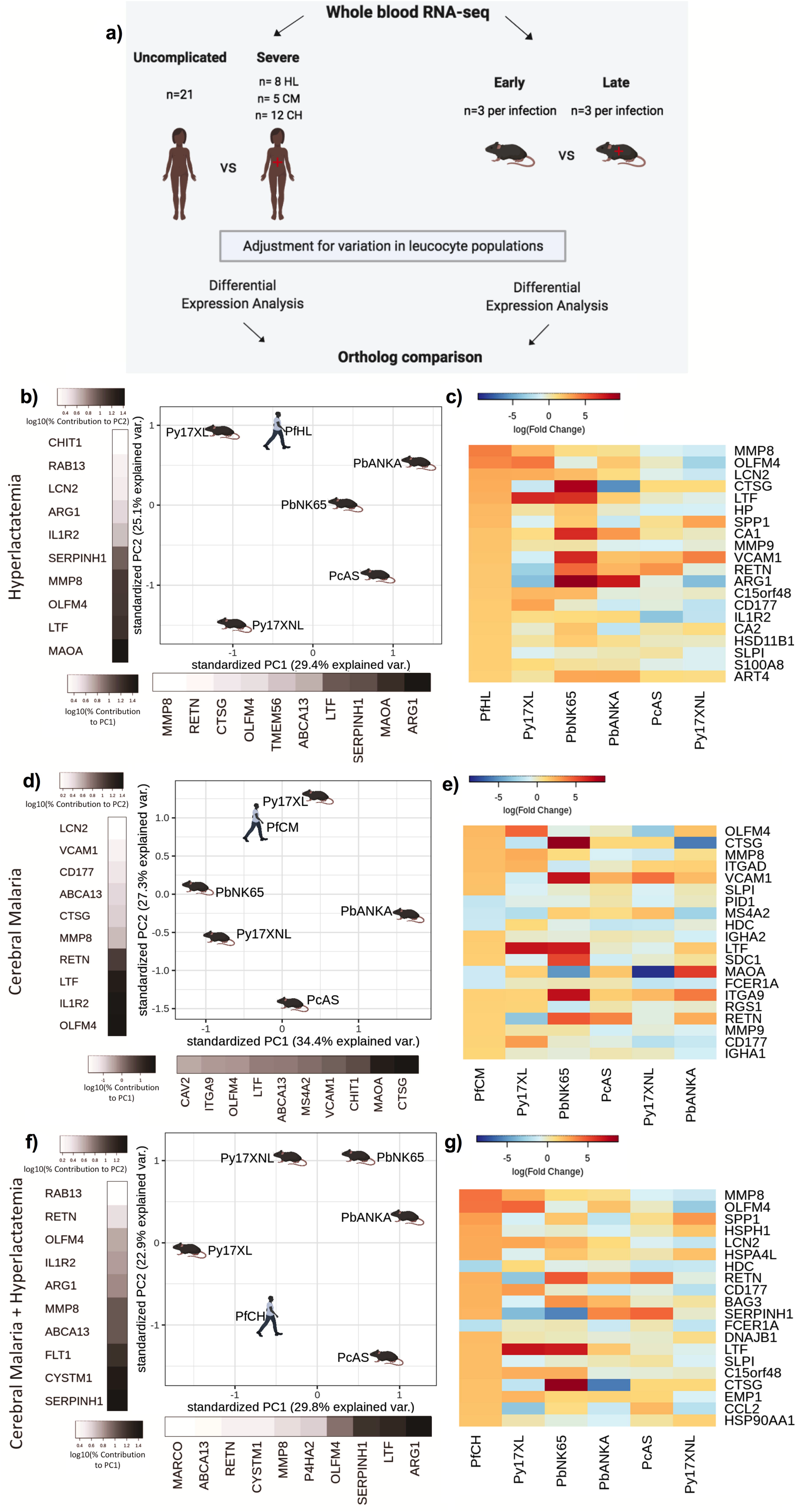
Comparison of host differential gene expression in three severe malaria phenotypes in Gambian Children and five mouse malaria models. a) Schematic illustration of the comparative transcriptomic analysis. b, d, f) Principal component analysis (PCA) plots generated using rank-normalised log fold change values from the human and mouse differential expression analyses. Only genes with 1: 1 mouse and human orthologs and with absolute logFC value greater than 1 in the corresponding human comparison were included. Comparison of changes in gene expression in the mouse models with those in human hyperlactatemia (PfHL) (b), cerebral malaria (PfCM) (d), or human hyperlactatemia plus cerebral malaria phenotype (PfCH) (f). The percentage of the total variation explained by principal components 1 and 2 are shown in the axis labels. Greyscale heatmaps parallel to each axis show the contributions of the 10 genes contributing most to the corresponding PC (c, e, g). Heatmaps show logFC for the 20 genes with the greatest absolute log fold change values in the human differential gene expression analysis, and their orthologs in each mouse model, corresponding to the analyses illustrated in b, d, and f, respectively. Mouse models are ordered left to right in order of increasing dissimilarity to the human disease, based on the Euclidian distance calculated from all principal components (Supplementary File 8). The rows (genes) are ordered by absolute log fold change in the human comparison in descending order. n=3 for early and n=3 for late time point in each mouse model; n= 21 Uncomplicated, n= 8 HL, n= 5 CM n= 12 CH. The mouse model abbreviations are as follows: PbNK65 (*P. berghei NK65*), PbANKA (*P. berghei ANKA*), PcAS (*P. chabaudi AS*), Py17XL (*P. yoelii 17XL*) and Py17XNL (*P. yoelii 17XNL*).

Overall, the direction of changes in gene expression in the mouse models were less concordant with those in human SM phenotypes than we observed in the comparisons with UM (Supplementary File 6, Fig. 3). There was, however, much clearer variation between the different mouse models in how closely the changes in expression of individual genes recapitulated those observed in each human SM manifestation (Fig. 3b, d, f). Using the principal component-based approach to compare weighted changes in gene expression in each infection, we were able to identify the models with greatest similarity to the transcriptional host response of each human SM phenotype (Fig. 3b, d, f and Supplementary File 6). It is notable that even amongst the twenty most differentially expressed genes associated with each human SM manifestation, there was considerable variation in the degree of concordance and discordance amongst the mouse models (Fig. 3c, e, g).

Hyperlactatemia is a relatively common manifestation of SM in children, and an independent predictor of death (21). Principal component analysis (PCA) revealed that *P. yoelii 17XL* and *P. berghei NK65* models most closely recapitulated the changes in gene expression associated with this disease phenotype in Gambian children (Fig. 3b). We performed gene ontology enrichment analysis on the genes contributing most to the principal components explaining the greatest proportion of variation between the mouse models and human disease, identifying neutrophil degranulation driving PC1 and myeloid leukocyte activation driving PC2 (Supplementary File 7). Despite *P. yoelii 17XL* having the closest proximity to human malaria hyperlactatemia in the PCA plot, it was clear that even for this model many of the most differentially expressed genes were not concordantly regulated (Fig. 3c, Supplementary File 6). Amongst the most concordant genes were those encoding neutrophil granule proteins: Lactotransferrin (*LTF, Ltf*), Olfactomedin 4 (*OLFM4, Olfm4*), CD177 molecule (CD177, *Cd177*), Matrix Metallopeptidase 8 (MMP8, *Mmp8*), Lipocalin 2 (LCN2, *Lcn2*), Matrix Metallopeptidase 8 (MMP9, *Mmp9*), and S100 Calcium Binding Protein A8 (S100A8, *S100a8*); but there was notable discordance of expression of genes encoding Arginase 1 (*ARG1, Arg1*), Cathepsin G (*CTSG, Ctsg*), Resistin (RETN, *Retn*), Vascular Cell Adhesion Molecule 1 (VCAM1, *Vcam1*), and Secreted Phosphoprotein 1 (SPP1, *Spp1*) (Fig. 3c).

In the comparison of the mouse models with the human CM phenotype (Fig. 3d), *P. yoelii 17XL* was again the mouse model with greatest similarity in gene expression changes, and gene ontology analysis revealed that myeloid leucocyte activation and neutrophil degranulation were again the most enriched GO terms amongst the genes explaining the greatest variation between models (Supplementary File 7). The genes with concordant and discordant changes in expression between humans and mice were also similar to those in the HL comparison.

Findings were similar when we compared the changes in gene expression in the mouse models with those in children with UM versus children with the most severe phenotype where both CM and HL are present (CH) (18). *P. yoelii 17XL* was placed closest to human CH in the PCA plot (Fig. 3g), and the genes contributing most to PC1 and PC2 were again enriched in neutrophil degranulation and myeloid leukocyte activation GO terms (Supplementary File 7). Taken together the comparisons between mouse models and these three SM phenotypes in Gambian children suggest that *P. yoelii* 17XL recapitulates the profile of the most prominent changes in gene expression associated with human SM phenotypes more closely than the other mouse models.

The relative frequency of different manifestations of *P. falciparum* SM varies across different geographic locations, influenced by the intensity of exposure to malaria, naturally acquired immunity, and age of the individual (22, 23). Changes in gene expression associated with the same disease manifestation may also vary between studies in different populations, under genetic and environmental influences, and due to technical differences in the methods used to assess gene expression (24, 25). Therefore, we investigated whether similar results would be obtained using data from an independent study conducted in Gabonese children with *P. falciparum* infection (20).

In the study from which we obtained this data, Gabonese children with CM and CH (CM/CH) were not distinguished as separate phenotypes and were pooled into a single group for microarray analysis (see methods). Nevertheless, comparison of changes in gene expression between early and late stages of the mouse infections with those between Gabonese children with UM and CM/CH revealed that *P. yoelii* 17XL most closely recapitulated the differential expression seen in humans (Fig. 4, Supplementary Files 2 & 4). Gene ontology analysis confirmed that the innate immune response and leukocyte mediated immunity were the main drivers of variation between models, similar to the analysis in Gambian children (Supplementary File 7).

**Figure 4:**
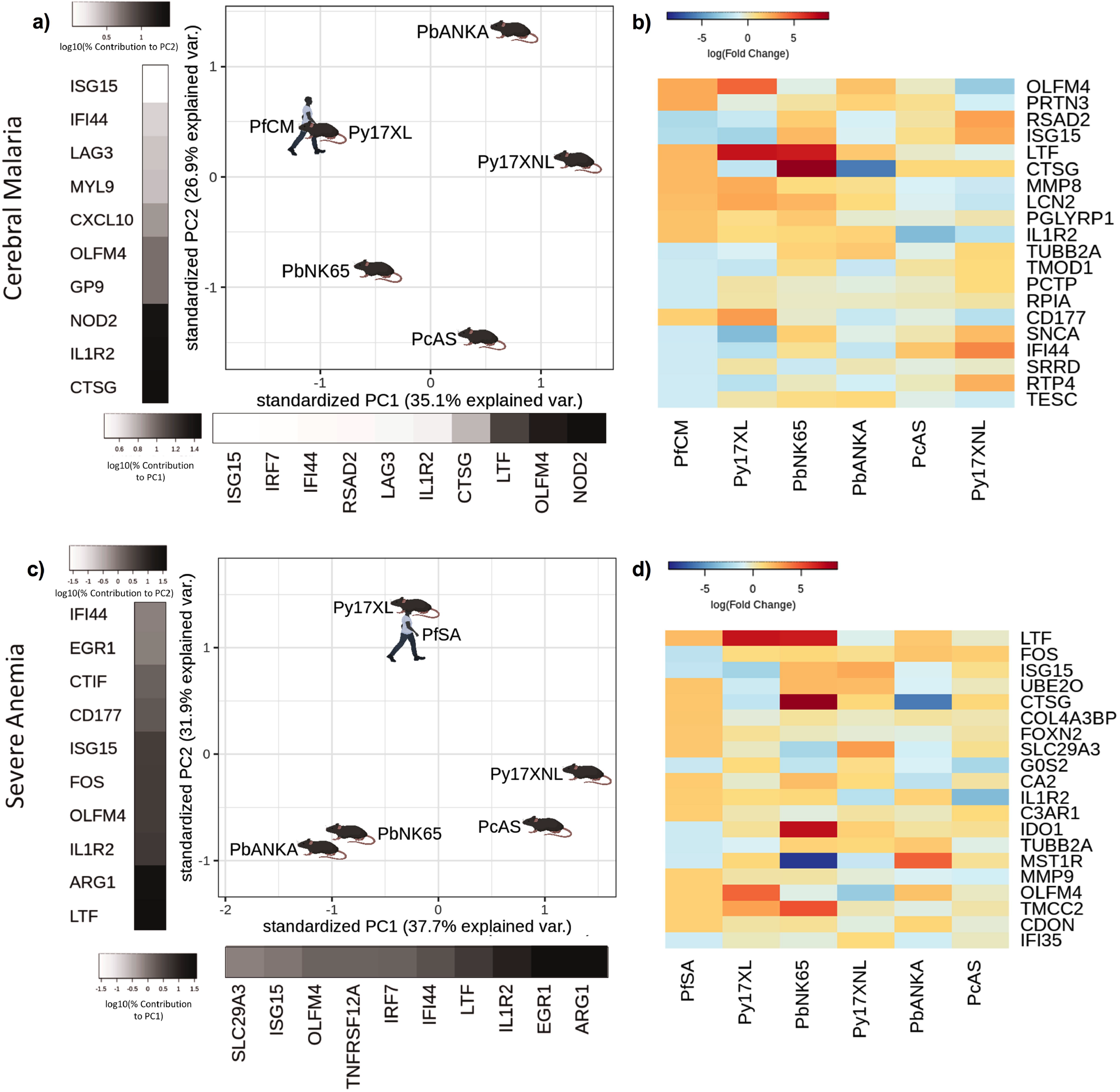
Comparison of host differential gene expression in two severe malaria phenotypes in Gabonese Children and five mouse malaria models. a, c) Principal component analysis (PCA) plots generated using rank-normalised log fold change values from the human and mouse differential expression analyses. Only genes with 1:1 mouse and human orthologs and with absolute logFC value greater than 1 in the corresponding human comparison were included. Comparison of changes in gene expression in the mouse models with those in human cerebral malaria (PfCM) (a) and severe anemia (PfSA) (c). The percentage of the total variation explained by principal components 1 and 2 are shown in the axis labels. Greyscale heatmaps parallel to each axis show the contributions of the 10 genes contributing most to the corresponding PC (b, d). Heatmaps show logFC for the 20 genes with the greatest absolute log fold change values in the human differential gene expression analysis, and their orthologs in each mouse model, corresponding to the analyses illustrated in a and c. Mouse models are ordered left to right in order of increasing dissimilarity to the human disease, based on the Euclidian distance calculated from all principal components (Supplementary File 8). The rows (genes) are ordered by absolute log fold change in the human comparison in descending order. n=3 for early and n=3 for late time point in each mouse model; n=5 pooled samples uncomplicated (UM), n=5 pooled samples CM, n=5 pooled samples SA (each pool contained RNA from 4 individuals with the same phenotype). The mouse model abbreviations are as follows: PbNK65 (*Plasmodium berghei NK65*), PbANKA (*Plasmodium berghei ANKA*), PcAS (*Plasmodium chabaudi AS*), Py17XL (*Plasmodium yoelii 17XL*) and Py17XNL (*Plasmodium yoelii 17XNL*).

In contrast to the Gambian dataset, where severe anemia (SA) was rare (26), the SA phenotype was included in the Gabonese dataset. Comparing the differential gene expression in the mouse models and those between UM and SA also identified that the changes in gene expression seen in *P. yoelii* 17XL were most similar to the differences seen in the Gabonese children (Fig. 4c). The genes with highly concordant expression between SA and *P. yoelii* 17XL were prominently neutrophil related (*LTF, OLFM4, MMP9* and *IL1R2*) (Fig. 4d), gene ontology analysis revealed that the main drivers of PC1 were slightly different to previous comparisons with prominence of immune response and type I interferon signalling pathways, whilst PC2 drivers were more similar to previous comparisons including leukocyte activation and neutrophil degranulation (Supplementary File 7). These data from Gabonese children provide independent, cross-platform, comparison, and substantiate that the profile of gene expression associated with severe *P. yoelii* 17XL infection is most similar to those in the major human SM phenotypes.

### Comparative transcriptomic results are consistent with pathophysiology

The profile of changes in gene expression associated with HL, CM, and SA, the three most common manifestations of SM in children, were all better recapitulated by the changes in gene expression in *P. yoelii* 17XL than any other mouse model. However, this model is not widely used to study the pathogenesis of these specific SM syndromes, so we sought to determine whether *P. yoelii* 17XL does reproduce the pathophysiological features of these infections. Blood lactate levels have rarely been reported in mouse malaria models, so we systematically measured lactate concentrations at early and late stages of infection in all five mouse models (Fig. 5a, Supplementary File 10). Small differences, if any, were noticed at the uncomplicated stage early in infection, while at maximum severity *P. yoelii* 17XL and *P. berghei* NK65 infected mice developed dramatic hyperlactatemia with concentrations similar to the maximum values seen in human HL (18).

**Figure 5.**
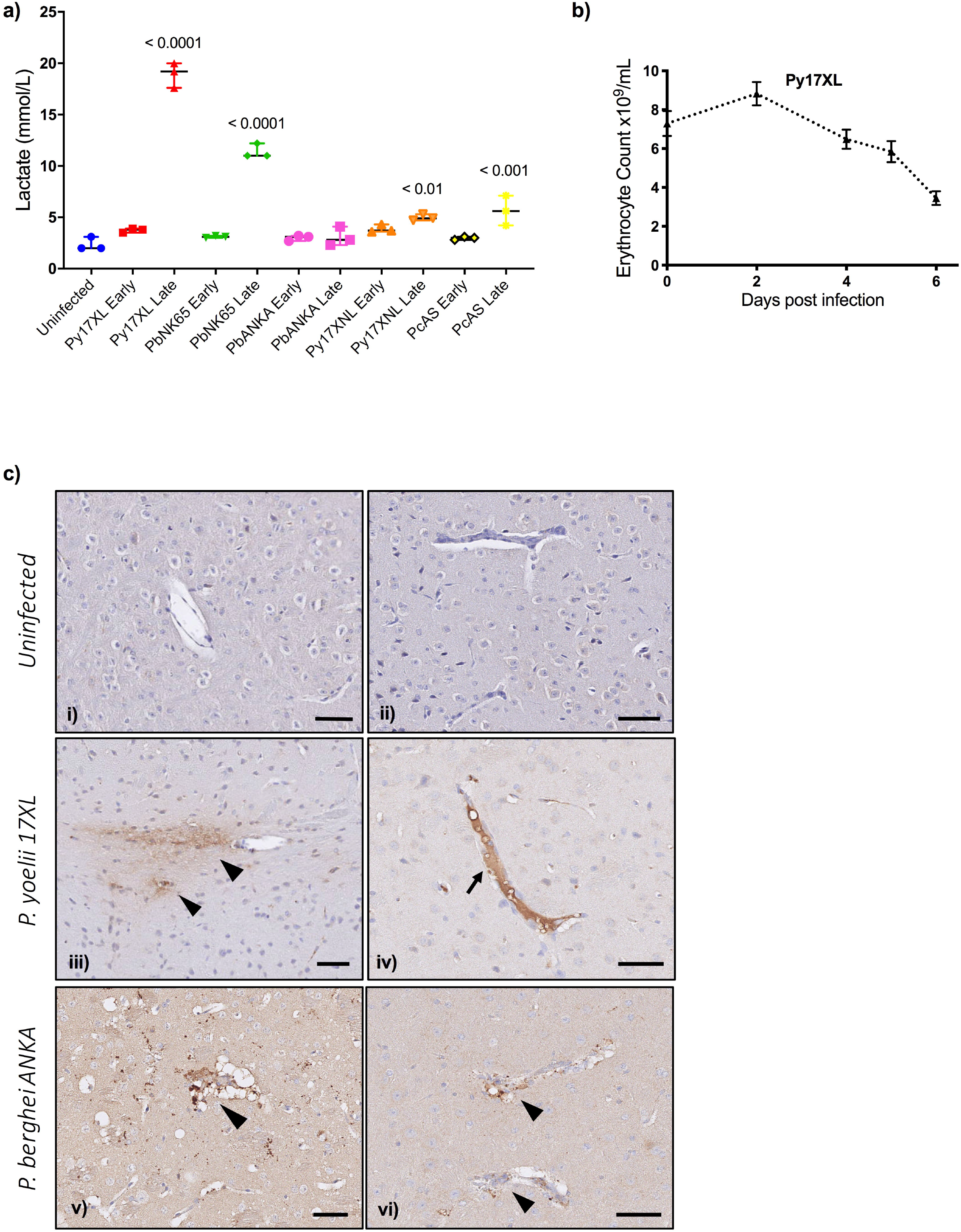
Pathophysiological features of rodent malaria infections. a) Lactate concentration in blood (mmol/L) in mice, uninfected, or at the early or late stage of each malaria parasite infection (n=3 for each infection time point). Error bars show median with range, One-way ANOVA *P-value* < 0.0001, *P-values* for post-hoc Dunnett’s multiple comparisons against uninfected mice are shown within the plot. b) Erythrocyte counts from *P. yoelii 17XL* infected mice, n=9, representative of 3 experiments, repeated measures ANOVA *P-value* < 0.01. c) Representative histological specimens of brain with fibrinogen staining to identify vascular leak in mice uninfected (i, ii), infected with *P. yoelii 17XL* (iii, iv), and infected *P. berghei ANKA* (v, vi) collected at the late stage (humane endpoint) of infection. Arrowheads identify extravascular fibrinogen indicating leak from the vasculature. Arrow points to strong intravascular fibrinogen staining (iv) suggestive of microthrombus. Representative images from analysis of uninfected mouse brains n=3; *P. yoelii* 17XL-infected mouse brains n=5; *P. berghei ANKA*-infected mouse brains n=4; Scale bar: 50 μm. 8-week-old wild-type female C57BL/6 mice were used in all experiments.

*P. yoelii* 17XL also reproduced the changes in gene expression associated with human SA better than other mouse models. Human SA is often associated with very high parasite biomass (26) and *P. yoelii* 17XL achieves much higher parasite load than other mouse models (Fig. 1) as well as causing rapid and profound anemia (27, 28) (Fig. 5b).

*P. yoelii* 17XL also showed the greatest transcriptional similarity to the pattern of changes in whole blood gene expression associated with human CM. *P. yoelii* 17XL was originally described as a virulent clone causing CM-like pathology (29), but it has subsequently been replaced by *P. berghei* ANKA as the most commonly used model of experimental CM. Since one of the key pathological mechanisms leading to death in paediatric CM is brain swelling due to extravascular fluid leak (30) we examined the presence of extravascular fibrinogen (31) as an indicator of vascular leak in the brains of both *P. berghei ANKA* and *P. yoelii 17XL* infected mice compared to uninfected (Fig. 5c). We found that brains from both infections had areas which stained positively for perivascular fibrinogen (indicative of vascular leak), while additionally some of the vessels from *P. yoelii 17XL* infected mice showed strong intravascular staining, suggestive of microthrombus formation (Fig. 5c iv), another mechanism that has been implicated in human CM (30, 31).

## Discussion

Mice are the most cost effective and widely used model organism for studying many human diseases (32, 33). Nevertheless, mice are distant evolutionarily and differ substantially from humans in many ways (5, 34). Disease models in mice often involve artificial induction of disease, which may reduce complexity and aid reproducibility, but might also limit their translational relevance. Therapeutic interventions that work in mice often fail when used in human clinical trials (35, 36). As a result, the usefulness of mice in some areas of translational research is debated (37, 38). Recently, concerted efforts have been made to improve both scientific and ethical aspects of the use of animals in biomedical research, with emphasis on the principles of replacement, reduction and refinement (the “3Rs”), and improving reproducibility through better experimental design and standardised reporting guidelines (39). Despite this, there has been little parallel effort made to assess or improve the relevance of animal models in translational research, and approaches which would improve translation from mice to humans are needed (40).

In malaria research mouse models are widely used but their relevance to human disease is contentious (8). Here, we objectively assessed the biological processes occurring in blood in some of the most commonly used mouse models of malaria to examine their similarity to human malaria, using a comparative transcriptomic approach. The five rodent malaria parasites we used led to the development of distinct disease trajectories and clinical features. Whilst no rodent malaria parasites induced changes in gene expression which fully recapitulated those in human malaria, at an early stage of infection, the rodent malaria parasites induced relatively similar transcriptional host responses to each other, with at least a broad overall similarity to that seen in a large study of UM in African children. However, when we investigated the similarity of the changes in gene expression associated with different SM manifestations, we saw that there was more heterogeneity, and the concordance and discordance of expression of individual genes varied more between each mouse model and each phenotype. One of the greatest sources of variation between the mouse models was in the myeloid cell response, particularly neutrophil response, associated with severe malaria manifestations.

An important implication of our findings is that the selection of the most appropriate mouse model for investigation of a particular mechanism of interest should not be made solely on the similarity of clinical phenotype in humans and mice. We propose that it should also be based on the degree of concordance of expression of genes associated with the mechanism of interest. Failure to consider the similarities and differences in biological processes indicated by gene expression could lead to experiments targeting pathways which are not involved in the host response to a particular mouse malaria parasite, making the experiments futile, unethical, and potentially leading to erroneous conclusions.

We identified that the pattern of changes in gene expression between early and late stages of *P. yoelii 17XL* infected mice showed the greatest similarity to the differences in gene expression between human UM and each of HL, CM, CH, and SA, suggesting that this model might be most representative of the profile of changes in host response induced by human severe malaria. This mouse model not only develops a very high parasite load, but our data suggest lethality at 5-7 days post infection is part of a multisystem disorder, accompanied by extreme hyperlactatemia at levels similar to those seen in human HL and CH. Until now, the lack of a rodent model to study malaria-induced hyperlactatemia has held back understanding of the mechanisms causing such high levels of lactate and how these relate to the increased risk of death in patients with malaria. *P. yoelii 17XL* infection of C57BL/6 mice is an attractive model for further translational research on this SM phenotype.

In the brains of *P. yoelii 17XL* infected mice we identified extravascular fibrinogen leak. This suggests that these mice may be in the process of developing a neurological syndrome at the time they reach the humane endpoint and may explain why this model showed transcriptional similarity to human CM. The transcriptional similarity of Py17XL to the human SA phenotype is consistent with the severe anemia and high parasite load which occurs in this infection.

Our study provides important insights into the translational relevance of commonly used mouse models of malaria, and more generally highlights the importance of considering relevance in addition to the 3Rs and reproducibility when planning any animal experiments. Our data is provided as a resource for researchers to help them to determine the concordance of gene expression between mouse malaria models and human disease, and we have identified an attractive mouse model for further translational studies on malarial hyperlactatemia. A strength of analysing the blood transcriptome is that it represents the systemic host response to infection, capturing both the direct influence of an infectious agent on blood leukocytes, and the response of blood leukocytes to mediators released into the circulation by cells in other organs. However, the blood transcriptome cannot assess the concordance of processes occurring within specific organs which do not produce changes in gene expression of circulating leukocytes, and our data should not be used to prevent testing of reasonable hypotheses about such tissue-specific interactions. Reassuringly, our findings were broadly consistent when we performed comparisons with human subjects from independent studies in different populations and using different transcriptomic methods. Stronger and more generalizable conclusions, and more nuanced approaches to analysis may be possible if future studies add to the data we have collected, with larger numbers of mice and greater sequencing depth. Future work should also assess other commonly-used mouse malaria models, using additional common mouse strains (including Balb/c and DBA/2), both sexes, and additional parasite strains.

## Methods

### Experimental design

We compared the whole blood transcriptome changes associated with severe malaria in mice and humans to identify concordant and discordant patterns of gene expression, and to identify which mouse models show the most similar changes to those seen in humans.

We chose to compare the changes in gene expression between human UM and SM categories with those seen between early and late mouse infections, assuming that mice harvested early in infection (when the first symptoms occur) represent uncomplicated malaria while mice at the peak of severity symptoms (or humane endpoints) represent severe malaria. Human data was obtained from published datasets from our group (18) and others (19, 20) while mouse data was generated specifically for this experiment.

### Animals and procedures

8-week-old wild-type female C57BL/6 mice were obtained from Charles River Laboratories. All mice were specified pathogen free, housed in groups of five in individually ventilated cages, and allowed free access to food and water. All protocols and procedures were approved by Imperial College Animal Welfare and Ethical Review Board, following Laboratory Animal Science Association good practice guidance. Mice were acclimatized to the animal facility for one week before any experimental procedures. Parasites kept in Alsever’s solution with 10% glycerol (mixed at 1:2 ratio) in liquid nitrogen were defrosted and accordingly diluted (depending on parasitemia of the frozen stock) to infect a passage mouse. The passage mouse infection was then closely monitored until healthy parasites were observed in a blood smear and parasitemia reached at least 2%. Sterile blood was collected, before parasitemia reached 5%, by cardiac puncture under non-recovery isoflurane anesthesia, and diluted in sterile PBS to achieve desired concentration. Viable parasites of strains *P. berghei ANKA* (lethal), *P. berghei NK65* (lethal), *P. yoelii 17XL* (lethal), *P. yoelii 17XNL* (non-lethal), and *P. chabaudi AS* (non-lethal) were prepared from frozen stocks by blood passage. Experimental mice were infected with 10^5^ live parasites by intraperitoneal injection. 50 mice were randomly allocated to be infected in groups of 10 with each parasite strain and then segregated into two cages of five mice each per parasite strain. 10 control uninfected mice were used for weight-gain comparisons.

The weight and physical condition of each mouse was monitored throughout the course of each infection (Supplementary File 11). Change in weight was calculated as a percentage of baseline weight measured prior to infection. For *P. berghei* ANKA infection, which causes experimental cerebral malaria, additional neurological monitoring was performed using the Rapid Murine Coma and Behaviour Scale (RMCBS) (13), which includes assessment of gait, motor performance, balance, limb strength, body position, touch escape, pina reflex, foot withdrawal reflex, aggression, and grooming. Due to the need for different intensity and nature of monitoring in each infection to ensure animal welfare, blinding to infection group was considered inappropriate.

The early time point was defined as the first time at which mice manifested any signs of ill health, including any reduction in activity, ruffled fur or weight loss. The late time point was defined as the humane endpoint for each lethal parasite species (Supplementary File 11), or a time point chosen to be just before the expected day of maximum severity of non-lethal infections (to avoid sampling mice which were starting to recover).

Tail capillary blood was used to prepare blood smears for analysis of parasitemia and lactate measurement using the Lactate Pro 2 (HAB direct) lactate meter. Parasitemia was quantified by microscopy of thin blood smears stained with 10% Giemsa and examined at 100x magnification with a Miller Square reticle. Erythrocyte counts were determined using a Z2 Coulter particle counter (Beckman Coulter). When mice were euthanized, heparinized blood was collected by cardiac puncture under non-recovery isoflurane anesthesia, and an aliquot of 300-500uL was immediately mixed at 1:2.76 volume ratio with fluid from a Paxgene blood RNA tube (Qiagen), whilst the remainder was stored on ice for flow cytometry analysis. Brains were collected from *P. yoelii* 17XL and *P. berghei* ANKA infected mice and fixed in 4% paraformaldehyde for 48 hours before being processed. Brains were then paraffin embedded, cut, and stained with antibody against fibrinogen (ab34269 1:100, Abcam, UK) using a Roche automated staining system. Digitized images taken at x 40 magnification (LEICA SCN400, Leica microsystems U.K) at IQPath (Institute of Neurology, University College, London, UK). Images were then viewed and examined with Aperio ImageScope software (Leica Biosystems Imaging, Inc.).

### Flow cytometry

The proportions of major leukocyte subpopulations in mouse blood were determined by flow cytometry using specific cell-surface marker antibodies. Approximately 50 μl of whole blood was mixed with 2 ml ammonium chloride red-cell lysis buffer for 5 minutes at room temperature, then samples were centrifuged and washed in flow cytometry buffer (specify this) and centrifuged again. Resultant cell pellets were resuspended in 50ul of antibody cocktail (all antibodies from Biolegend, Supplementary File 12) for 30 minutes before further washing and fixation in 2% paraformaldehyde. Flow cytometry was performed using a BD LSR Fortessa machine. BD FACSDiva software was used to collect the data and analysis was conducted using FlowJo v10 (TreeStar Inc.), gating on single leukocytes before identification of major cell populations according to their surface marker staining (Supplementary Figure 1 in Supplementary File 1). Leucocyte proportions for early and late timepoint within each infection are presented in Supplementary Figure 2 in Supplementary File 1.

### RNA isolation from mouse blood

RNA extraction was performed using the PAXgene Blood RNA Kit (Qiagen) according to the manufacturers’ instructions (41). After the isolation of the RNA, Nanodrop ND-1000 Spectrophotometer (Labtech) was used to obtain the ratio of absorbance at 260 nm and 280 nm (260/280) which is used to assess the purity of RNA (or DNA). Values of ~2 are generally accepted as pure for RNA. RNA integrity was assessed using Agilent RNA 6000 Nano Kit (Agilent), used according to the manufacturers’ instructions with the Agilent 2100 Bioanalyzer (Agilent), and all traces were inspected visually for evidence of RNA degradation because the RNA Integrity Number calculation can be misleading when host and parasite RNA are both present in significant quantities (18).

For the RNA sequencing analysis 6 samples were selected from each infection (3 from the early time point and 3 from the late time point), along with 3 uninfected control. Samples were selected based on the RNA quality (260/280 ratio and Agilent 2100 Bioanalyzer traces). If more than 3 samples for each infection and time point were of sufficient quality, we selected the three with most similar clinical score and parasitemia levels within each group.

### Dual-RNA sequencing

Library preparation and sequencing to generate the mouse RNA-Seq data was performed at the Exeter University sequencing service. Libraries were prepared from 1μg of total RNA with the use of ScriptSeq v2 RNA-Seq library preparation kit (Illumina) and the Globin-Zero Gold kit (Epicentre) to remove globin mRNA and ribosomal RNA. Prepared strand-specific libraries were sequenced using the 2×125bp protocol on an Illumina HiSeq 2500 instrument.

### Gene annotations

Human reference genome (hg38) was obtained from UCSC genome browser (http://genome.ucsc.edu/), mouse reference genome (mm10) was obtained from UCSC genome browser (http://genome.ucsc.edu/). Human gene annotation was obtained from GENCODE (release 22) (http://gencodegenes.org/releases/), mouse gene annotation was obtained from GENCODE (release M16) (http://gencodegenes.org/releases/). The *Plasmodium* (*P. berghei, P. chabaudi and P. yoelii*) genomes were obtained from PlasmoDB (release 24) (42).

### Mouse RNA-Seq Quality Control, Mapping and Quantification

Quality control was carried out using fastqc (43) and fastqscreen (44). Adapters were trimmed using cutadapt (45). The read 1 (R1, -a) adapter is AGATCGGAAGAGCACACGTCT, and the read 2 (R2, -A) adapter is AGATCGGAAGAGCGTCGTGTAGGGAAAGAGTGT.

The trimmed reads were then mapped to the combined genomic index containing both mouse and the appropriate *Plasmodium* genome using the splice-aware STAR aligner (46). Reads were extracted from the output BAM file to separate parasite-mapped reads from mouse-mapped reads. Reads mapping to both genomes were counted for each sample and removed. BAM files were sorted, read groups replaced with a single new read group and all reads assigned to it. HTSeq-count (47) was used to count the reads mapped to exons with the parameter “-m union”. Only uniquely mapping reads were counted.

### Mouse Differential gene expression analysis

The raw expression counts can be found in Supplementary File 13. Firstly, the ensembl gene ID versions were matched to their MGI gene symbols and entrez IDs using biomaRt (annotation used: http://jul2018.archive.ensembl.org, mmusculus_gene_ensembl) (48, 49). Genes for which this information was not available were excluded from the analysis. Of these, only genes with raw expression values of greater than 5 in at least 3 samples were taken forward.

The differential gene expression analysis was then performed using the R package edgeR, raw read counts of each data set were normalised using a trimmed mean of M-values (TMM), which considers the library size and the RNA composition of the input data.

In order to account for variation between samples in the proportions of the major blood leukocyte populations (neutrophil, monocyte, CD4 T cell, CD8 T cell) we used their proportions estimated by flow cytometry (Supplementary File 14) as covariates in edgeR, adjusting for their effect on whole blood gene expression. B cells were excluded from the design matrix of the differential expression analysis due to the proportions totalling 100%. Thus, the design matrix (with the intercept set to 0) consisted of each sample’s disease type (the mouse model plus if the sample was early or late in infection i.e. *P. yoelii 17XL*_late) with the cell type proportions as covariates. Results of the differential expression analyses are presented in Supplementary File 2. Metadata matches each sample to their phenotype can be found in Supplementary File 15.

### Analysis of the Human RNA-Seq dataset

For the comparison with RNA-Seq data from human hosts, data from our previously published Gambian children cohort (Supplementary File 1) were used (18). This dataset can be found in the ArrayExpress database (www.ebi.ac.uk/arrayexpress) using the accession number E-MTAB-6413 and metadata are also presented in Supplementary File 16. The differential expression results were extracted from this study. These had already been adjusted for variation in leukocyte proportions and were used without further processing. Lists of differentially expressed genes in Supplementary File 9.

### Analysis of Human Microarray Datasets

Expression values for two human microarray datasets were extracted from the GEO database (20, 50). For the Boldt et al. study, background correction, normalisation, and batch correction was performed on the raw expression values using the methods given in Supplementary File 1. For the Idaghdour et al. study the data was downloaded as pre-normalised expression values.

CellCODE (51) was used to estimate the proportions of the major blood leukocyte subpopulations (neutrophils, monocytes, CD4 T cells, CD8 T cells and B cells) in each of the samples, based on reference gene expression profiles (Allantaz et al. GEO Accession: GSE28490 (52); the full signature dataset derived from Allantaz et al., not just those used for these datasets, can be found in Supplementary File 17). Surrogate proportion variables for each leukocyte subpopulation were then used as covariates in differential gene expression pairwise analysis in Limma (53) (Supplementary File 1).

The One sample (GSM848487) was removed from the Idaghdour et al. dataset because the age of the subject was not available. The original study sampled a population with wide age range from different locations, so following the approach in the original study, differential expression analysis included age, location (Zinvie or Cotonou), and hemoglobin genotype (AA, AS or AC) in addition to the leukocyte subpopulation surrogate proportion variables as covariates for the pairwise differential expression analysis conducted using Limma.

The lists of differentially expressed genes for these datasets are available in Supplementary Files 3 and 4.

### Identification of Orthologous Genes

A text file of all the orthologous (Ensembl 52) *Homo sapiens* (NCBI36) and *Mus Musculus* genes was extracted from the Ensembl database and used as a reference (Supplementary File 5).

### Comparative Transcriptomics using Principal Component Analysis

To use as much information as possible about changes in gene expression between conditions in human and mouse malaria datasets of varying size, we did not impose a p-value threshold but began by selecting all genes in the human differential expression analyses with absolute log-fold change greater than 1. We then selected those with 1:1 orthologs in mice, and used these genes for subsequent comparisons with gene expression in mice. There were no cut-offs applied based on the differences in expression between early and late stage infection in mice. Therefore, our analyses assess the extent to which changes in mouse gene expression recapitulate those in humans, but do not address the reciprocal question of how well human gene expression recapitulates that in mice.

To compare patterns of gene expression associated with pathogenesis between species, without undue influence of species-specific variation in the baseline- or inducible-expression of each gene, we focused further analysis on the contrasts between comparable pairs of human and pairs of mouse infection states. Both human microarray UM vs healthy results were compared to the mouse early stage infection vs uninfected control results.

The human RNA-Seq (Lee et al. 2018) CM vs UM, HL vs UM, CH vs UM, and microarray (Boldt et al. data) CM vs UM and SA vs UM results were compared to mouse late stage vs early stage of infection results for each mouse model.

To allow comparison of the relative magnitudes of changes in gene expression between the human and mouse models, we developed a rank-based analysis of the changes in expression in each human and mouse pairwise comparison. Genes were ranked in descending order of absolute log fold change, with ties given the same minimum rank. Each gene was then assigned a value of 100 divided by rank, which was then multiplied by the sign of the original log fold change. For example, if the original log fold change was negative the rank-standardised value would then be multiplied by −1. This approach means that the genes with greatest difference in expression between the conditions of interest within-species have the biggest effect on the comparative transcriptomic analysis between species. These values are presented in Supplementary File 18 and were used as the input for subsequent Principal Component Analysis (PCA) to highlight the differences and similarities between the mouse models and human disease comparisons in low-dimensional space. The PCAs were performed using the R-core function Prcomp() with default parameters and visualised using functions from the ggbiplot (54) and ggimage packages. The 10 genes that contributed the most to principal components 1 and 2 (a subset of those given in Supplementary File 19) were collected using the factoextra (55) and FactoMineR (56) packages, specifically the PCA() function, with scale.unit set to FALSE to correspond to the default parameters of the Prcomp() function.

### Gene Ontology Analysis

Lists of genes contributing greater than or equal to 0.1% to PC1 and/or PC2 were also extracted (Supplementary File 19). These were used as the genes of interest for Gene Ontology (GO) term enrichment analysis performed using the goana.DGELRT() function (Package: Limma) (53). The list of all the 1:1 orthologs used as the input for the Principal Component Analysis were used as the background gene lists (Supplementary File 20). Human gene IDs were fed to the GO term enrichment analysis. For each comparison in each dataset the Euclidean distances (Supplementary File 8) between each of the mouse models and the human data were calculated using standardised log fold change values and the R-core dist() function.

### Heatmaps

The 20 genes with the greatest absolute log fold change value in the each human disease comparison were used to construct illustrative heatmaps using the heatmap.2() function from the R package gplots (57).

### Statistical Tests

GraphPad Prism 8 (GraphPad Software) was used for statistical analyses of lactate concentration in the different mouse models and erythrocyte counts from *P. yoelii 17XL* infected mice. One-way ANOVA test was used to compare the lactate concentration in mice uninfected or infected at different time points and post-hoc Dunnett’s test for multiple comparisons. One-way ANOVA for repeated measures was used to analyse erythrocyte counts from *P. yoelii 17XL* infected mice. All tests were two-sided using a significance threshold of 5%.

## Supporting information

Supplementary File 1 Supplementary Figures and small tables

Supplementary File 2 Mouse Differential Expression Analysis

Supplementary File 3 Idaghdour et al 2012 Differential Expression Analysis

Supplementary File 4 Boldt et al 2019 Differential Expression Analysis

Supplementary File 5 Human Mouse Orthologs

Supplementary File 6 Discordance Concordance

Supplementary File 7 GO terms

Supplementary File 8 Euclidean Distances

Supplementary File 9 Lee et al 2018 Differential Expression Analysis

Supplementary File 10 Mouse Lactate Measurements

Supplementary File 11_severity scoring

Supplementary File 12 Antibodies used for FACS

Supplementary File 13 RNA-Seq Mouse Raw Counts

Supplementary File 14 Mouse Cell Type Proportions

Supplementary File 15 Metadata Sample IDs Phenotypes

Supplementary File 16 Lee at al 2018 Sample Metadata

Supplementary File 17 Reference Immune Cell Type Expression Profile for Boldt and Idaghdour

Supplementary File 18 PCA input standardised logFC values

Supplementary File 19 Genes contributing to PC1 and PC2

Supplementary File 20 GO Term Background Gene Lists

## Acknowledgements

This work was supported by the UK MRC and the UK Department for International Development (DFID) under the MRC/DFID Concordat agreement and is also part of the EDCTP2 program supported by the European Union (MR/L006529/1 to A.J.C.), Imperial College Dean’s EPSRC Studentship (to C.D.), Imperial College-Wellcome Trust Institutional Strategic Support Fund (to A.G.), Sir Henry Wellcome Fellowship (206508/Z/17/Z to M.K), and the analysis of patient data was supported by the NIHR Imperial Biomedical Research Centre. Exeter Sequencing Service is supported by an MRC Clinical Infrastructure award (MR/M008924/1), the Wellcome Trust Institutional Strategic Support Fund (WT097835MF), a Wellcome Trust Multi-User Equipment Award (WT101650MA), and a Biotechnology and Biological Sciences Research Council Longer and Larger award (BB/K003240/1).

## Author contributions

A.G. and A.J.C. contributed to conceptualization; A.G. performed experimental work; A.G., C.D., P.S.B., and H.J.L. analyzed the data; A.G., C.D., A.J.C. and M.K. drafted the manuscript; A.J.C. supervised all aspects of the work; all authors contributed to critical revision and approved the final manuscript.

## Materials & Correspondence

Correspondence and requests for access to data and materials should be addressed to Dr Aubrey J. Cunnington.

## Data Availability

The adapter trimmed RNA-Sequencing files for the mouse RNA-Seq data have been submitted to the European Nucleotide Archive (ENA) and are available under the study accession number PRJEB43641.

## Supplementary Materials

Supplementary File 1 Supplementary Figures and small tables

Contains:

Supplementary Table. 1 Details of the publicly available human datasets

Supplementary Figure 1: Gating strategy for defining WBC proportions in mouse blood

Supplementary Figure 2: Leucocyte proportions measured in whole blood by flow cytometry.

Supplementary File 2 Mouse Differential Expression Analysis

Supplementary File 3 Idaghdour et al 2012 Differential Expression Analysis

Supplementary File 4 Boldt et al 2019 Differential Expression Analysis

Supplementary File 5 Human Mouse Orthologs

Supplementary File 6 Discordance-Concordance Table

Supplementary File 7 Gene ontology enrichment analysis

Supplementary File 8 Euclidean Distances

Supplementary File 9 Lee et al 2018 Differential Expression Analysis

Supplementary File 10 Mouse Lactate Measurements

Supplementary File 11 Severity scoring

Supplementary File 12 Antibodies used for FACS

Supplementary File 13 Mouse RNA-Seq Raw Counts

Supplementary File 14 Mouse Cell Type Proportions

Supplementary File 15 Metadata for Mouse RNA-Seq Files

Supplementary File 16 Lee at al. 2018 Sample Metadata

Supplementary File 17 Reference Immune Cell Type Expression Profiles for Deconvolution

Supplementary File 18 PCA input standardised logFC values

Supplementary File 19 Genes contributing to Principal Component Analyses

Supplementary File 20 GO Term Background Gene Lists

## Supplementary figure legends

**Supplementary Figure 1: Gating strategy for defining WBC proportions in mouse blood**

The strategy included gating around the WBC population excluding red blood cells that did not lyse and debris using FSC-A/SSC-A. Then doublets were excluded with FSC-A and FSC-H. Using different combinations of antibodies or antibody/ SSC-A proportions of the populations of interest were defined. T cells gating: CD3 +, CD19-; T helper cells: CD8a-, CD4+; cytotoxic T cells: CD8a+, CD4-; B cells: CD3-, CD19+; Monocytes: CD11b +, Ly-6G-; Neutrophils: Ly-6G+, CD11b +.

**Supplementary Figure 2: Leucocyte proportions measured in whole blood by flow cytometry.**

8-week-old female wild type C57BL/6 mice infected with: *P. yoelii 17XL, P. berghei ANKA, P. berghei NK65, P. yoelii 17XNL, P. chabaudi AS*, and uninfected controls are presented here. Proportions of B cells, monocytes, neutrophils, T helper cells and cytotoxic T cells were measured at the early and late time point of each infection and compared to uninfected mice. n=3 for early and n=3 for late time point in each mouse model; n=3 for uninfected mice. Bars show mean with 95% CI. The mouse model abbreviations are as follows: PbNK65 (*P. berghei NK65*), PbANKA (*P. berghei ANKA*), PcAS (*P. chabaudi AS*), Py17XL (*P. yoelii 17XL*) and Py17XNL (*P. yoelii 17XNL*).

