## Supplementary File 1 Supplementary Figures and small tables for "Comparative transcriptomic analysis reveals translationally relevant processes in mouse models of malaria"

### Supplementary Table.1 Details of the publicly available human datasets

| Accession Number | Citation | Clinical Phenotypes | Numbers of subjects | Gene Expression Platform | Background correction Method | Normalisation Method | Differential Expression Analysis Comparison(s) |
|---|---|---|---|---|---|---|---|
| GEO: GSE1124 | Boldt et al. 2019 | Healthy<br><br>Severe Anaemia (SA)<br><br>Cerebral Malaria (CM)<br><br>Uncomplicated Malaria (UM) | Healthy (n=5 pools of 4 patients), SA (n=5 pools of 4 patients), CM (n=5 pools of 4 patients), UM (n=5 pools of 4 patients) | Microarray<br><br>Affymetrix Human Genome U133A Array | normexp | Quantile Normalisation | CM-Healthy<br><br>SMA-Healthy<br><br>UM-Healthy |
| GEO: GSE34404 | Idaghdour et al. 2012 | Uncomplicated Malaria (UM)<br><br>Healthy | UM (n=93), Healthy (n=61)<br><br>1 symptomatic sample was removed because its age information was not available | Microarray<br><br>Illumina HumanHT-12 V4 Expression BeadChip | Imported as background corrected values. In the original paper background values were subtracted using the averaging of the negative control probes | Imported as normalised values, the original paper performed quantile normalisation | UM -Healthy |
| ArrayExpress: E-MTAB-6413 | Lee et al. 2018 | Uncomplicated Malaria (UM)<br><br>Cerebral Malaria (CM)<br><br>Hyperlactatemia (HL)<br><br>Cerebral Malaria and Hyperlactatemia (CH) | UM (n=50)<br><br>CM (n=14)<br><br>HL (n=16)<br><br>CH (n=26) | RNA-Seq Illumina HiSeq 2500 | Not applicable | Imported the results of the differential expression analysis. The expression values were normalised by the originally paper using the trimmed mean of M-values method | HL-UM<br><br>CM-UM<br><br>CH– UM |

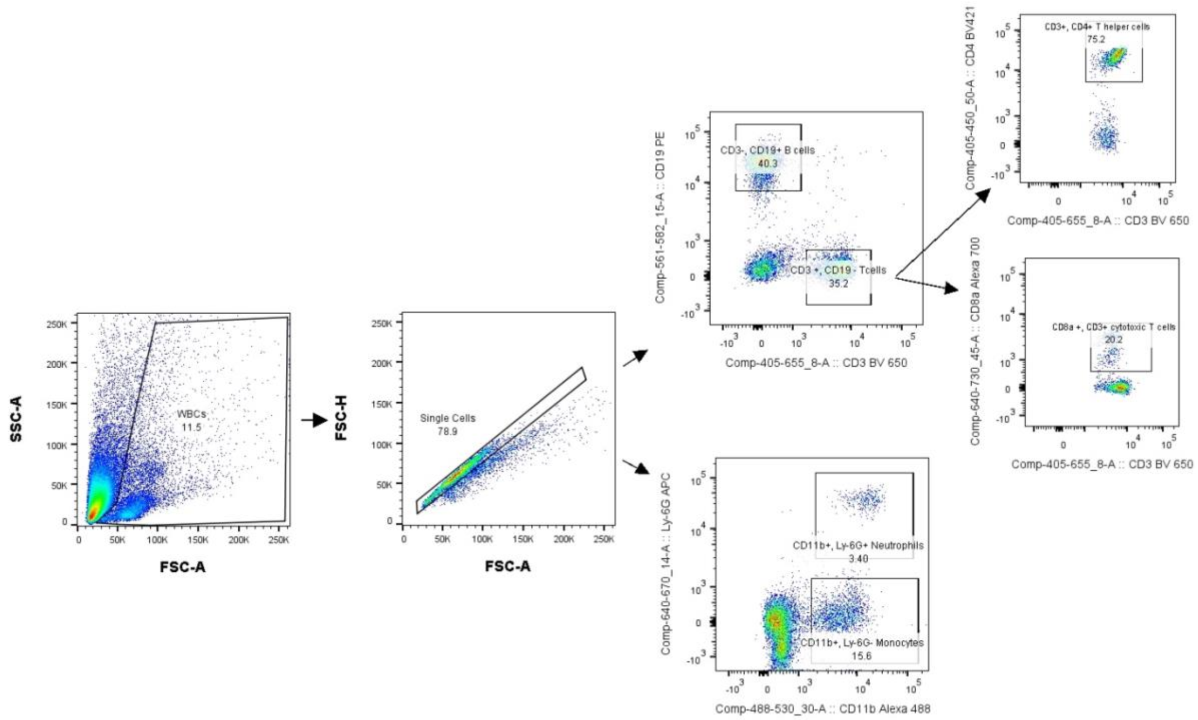


**Supplementary Figure 1: Gating strategy for defining WBC proportions in mouse blood**

The strategy included gating around the WBC population excluding red blood cells that did not lyse and debris using FSC-A/SSC-A. Then doublets were excluded with FSC-A and FSC-H. Using different combinations of antibodies or antibody/ SSC-A proportions of the populations of interest were defined. T cells gating: CD3 +, CD19-; T helper cells: CD8a- , CD4+; cytotoxic T cells: CD8a+, CD4-; B cells: CD3-, CD19+; Monocytes: CD11b +, Ly-6G-; Neutrophils: Ly-6G+, CD11b +.

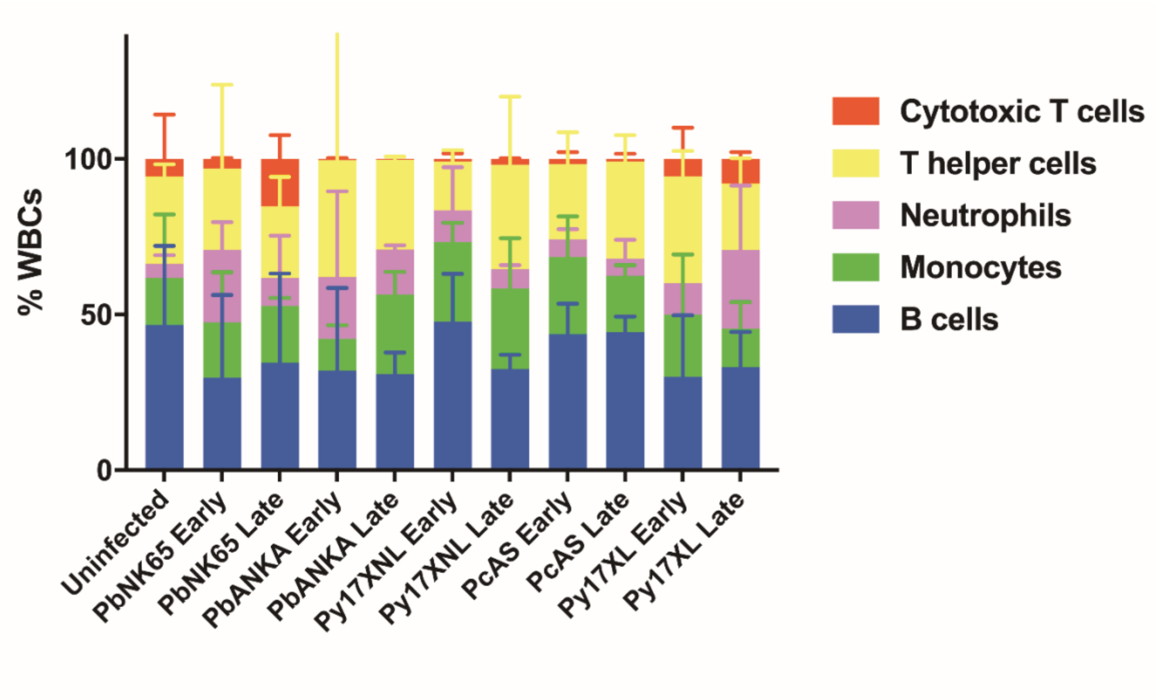


**Supplementary Figure 2: Leucocyte proportions measured in whole blood by flow cytometry.**

8-week-old female wild type C57BL/6 mice infected with: *P. yoelii 17XL*, *P. berghei ANKA*, *P. berghei NK65*, *P. yoelii 17XNL*, *P. chabaudi AS*, and uninfected controls are presented here. Proportions of B cells, monocytes, neutrophils, T helper cells and cytotoxic T cells were measured at the early and late time point of each infection and compared to uninfected mice. n=3 for early and n=3 for late time point in each mouse model; n=3 for uninfected mice. Bars show mean with 95% CI. The mouse model abbreviations are as follows: PbNK65 (*P. berghei NK65*), PbANKA (*P. berghei ANKA*), PcAS (*P. chabaudi AS*), Py17XL (*P. yoelii 17XL*) and Py17XNL (*P. yoelii 17XNL*).
