## Supplementary File 11_severity scoring for "Comparative transcriptomic analysis reveals translationally relevant processes in mouse models of malaria"

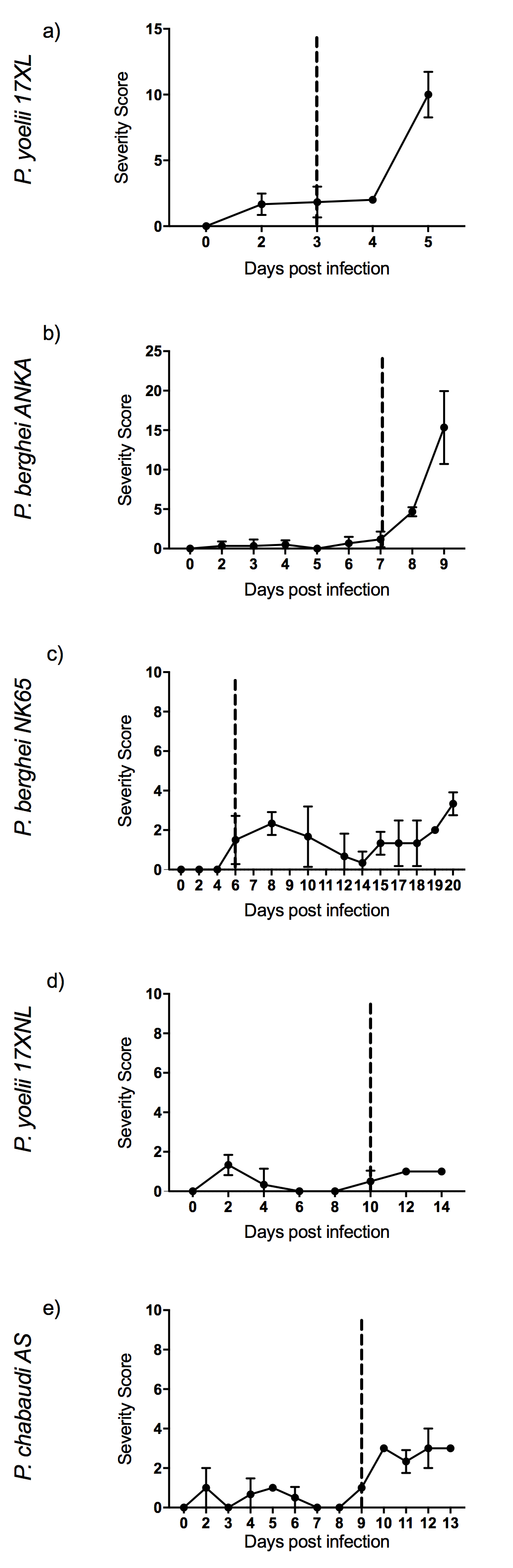

**Severity scoring in five mouse malaria models:** 8-week-old C57BL/6 female wild type mice infected with: *P. yoelii 17XL, P. berghei ANKA, P. berghei NK65*, *P. yoelii 17XNL*, and *P. chabaudi AS, respectively*. Points show mean, and bars show SD, for n=6 mice (up to and including time point of first signs of ill health, dashed vertical line) and n=3 mice (after dashed vertical line) for each infection.

SCORE SHEET FOR SCORING ENDPOINTS IN RODENTS

FOR USE WITH PROTOCOL 1 ONLY (Pathogen Maintenance) – MODERATE SEVERITY LIMIT

**PPL details number and Group:**	PPL 70/8919, Systemic Host Pathogen Interactions And Disease, A. Cunnington

**Date: Time**:	**PIL Number:**	**Name of person scoring**: **Name of Supervisor/Chief Investigator: (PPL or Supervisor):** **Contact Telephone Number:**	**After Hours:**

| Indicators | Scoring of independent variables: | Animal Number and Score |
| --- | --- | --- |
| **Date** |  |  |
| **General Health**  Eating  Locomotion  Behaviour: | 0. drinking and eating well 1. change in eating or drinking habit 2. reduction in skin elasticity 3. not eating/drinking, severely dehydrated 0. walking normally 1. limping, stiffness 2. laboured , ataxia 3. severely restricted mobility, paralysis of any limb 0. normal 1. away from littermates 2. aggressive or huddled in corner 3. severe distress, seizures |  |
| Appearance | 0. Normal  1. ruffled fur  2. animal appears depressed , hunched, reluctant to move  3. animal appears severely depressed, severe respiratory distress |  |
| **Weight loss*** | 0. normal 1. 5 – 10% 2. 10-19% (or body condition score <2) 3. >20% |  |
| Total Score |  |  |

###### For Total Scores

0 = normal: continue to monitor according to protocol	***** *A score of 3 in any one category: euthanase / Schedule 1 kill***

1-2 = mild changes: should be monitored at least once daily >2 = euthanase

* Assessment of Weight: Weight should be measured at the same time every other day. For experiments lasting up to 5 days, weight loss is calculated relative to the body weight immediately prior to start of the infection (or procedure if uninfected). For experiments lasting more than 5 days, this calculation must include adjustment for the expected weight gain that would occur in normal mice – calculated from the average weight gain of healthy control mice of the same age and gender as the experimental animals. Body condition score is determined according to the method of Ullman-Culleré and Foltz, 1999

**Signature of person scoring:…………………………………………………………………..**

**1. Morton, D.B. (1997). A scheme for the recognition and assessment of adverseeffects. In:Animal Alternatives, Welfare and Ethics. (Eds. LFM vanZutphen and M. Balls). ElsevierScience, Amsterdam. pp. 235-241.**

**2. Morton, D.B. (1998). The use of score sheets in the implementation of humaneend points.Proceedings of the Joint ANZCCART / NAEAC Conference on EthicalApproaches to Animalbased Science (Eds. D. Mellor, M. Fisher and G.Sutherland), ANZCCART, Adelaide andWellington. pp: 75-82.**

**EXAMPLE SCORE SHEET FOR SCORING ENDPOINTS IN RODENTS**

**FOR USE WITH PROTOCOL 2 ONLY (*Determinants of severe malaria*) – SEVERE SEVERITY LIMIT**

**PPL details number and Group:** PPL 70/8919, Systemic Host Pathogen Interactions And Disease, A. Cunnington

**Date:	Time**:	**PIL Number:**	**Name of person scoring**: **Name of Supervisor/Chief Investigator: (PPL or Supervisor):**

**Contact Telephone Number:	After Hours:**

| **Indicators** | **Scoring of independent variables:** | **Animal Number and Score** |
| --- | --- | --- |
| **Date** |  |  |
| **Appearance** | **0 Normal** |  |
|  | **1 General lack of grooming** |  |
|  | **2 Coat staring, ocular and/or nasal discharges** |  |
|  | **3 Piloerection, hunched up** |  |
|  | **Baseline weight** |  |
|  | **Baseline lactate** |  |
|  | **Baseline glucose** |  |
| **Food and water intake** | **0 Normal** |  |
|  | **1 Uncertain: body weight* < 5% reduced** |  |
|  | **2 Reduced intake: body weight* 5-15% reduced or body condition score <2** |  |
|  | **3 No food or water intake** |  |
| **Natural behaviour** | **0 Normal** |  |
|  | **1 Minor changes** |  |
|  | **2 Less mobile and alert, isolated** |  |
|  | **3 Vocalisation, self-mutilation, restless or still** |  |
| **Experimental Cerebral malaria** | **10 Ataxia** |  |
|  | **Seizures or generalised paralysis**   - **Euthanise** |  |
| **Provoked behaviou**r | **0 Normal** |  |
|  | **1 Minor depression or exaggerated response** |  |
|  | **2 Moderate change in expected behaviour** |  |
|  | **3 Reacts violently or very weak and precomatose** |  |
| **Score** | **If you have scored a 3 more than once, score an extra point for each 3 (2-4)** |  |
|  | **Total (0-26)** |  |

**0-3 = Normal.**

**4-8 = Monitor carefully, consider supportive care if appropriate.**

**9-13 = Suffering, provide relief, observe at least twice daily regularly. Seek advice from NACWO and/or NVS.**

**14-15 = Monitor every 4 hours. Provide support as directed by NACWO and/or NVS**

**16+ = Euthanise animal**

* Assessment of Weight: Weight should be measured at least once daily, at the same time every day

For experiments lasting up to 5 days, weight loss is calculated relative to the body weight immediately prior to start of the infection (or procedure if uninfected)

For experiments lasting more than 5 days, this calculation must include adjustment for the expected weight gain that would occur in normal mice – calculated from the average weight gain of healthy control mice of the same age and gender as the experimental animals.

Body condition score is determined according to the method of Ullman-Culleré and Foltz, 1999

**Reference:**

**Practical use of distress scoring systems in the application of humane endpoints. M.H Lloyd & S.E. Wolfensohn**

SCORE SHEET FOR SCORING ENDPOINTS IN RODENTS

FOR USE WITH PROTOCOL 2 ONLY (Determinants of severe malaria) – SEVERE SEVERITY LIMIT

**PPL details number and Group:**	PPL 70/8919, Systemic Host Pathogen Interactions And Disease, A. Cunnington

**Date: Time**:	**PIL Number:**	**Name of person scoring**: **Name of Supervisor/Chief Investigator: (PPL or Supervisor): Contact Telephone Number:**

| **Indicators** | **Scoring of independent variables:** | **Animal Number and Score** |
| --- | --- | --- |
| **Date** |  |  |
|  | **Baseline weight (g)** |  |
|  | **Blood lactate (mmol/L, if measured)** |  |
|  | **Blood glucose (mmol/L, if measured)** |  |
| **Body weight and condition** | **10 Normal (baseline +/- 0.2g)** |  |
|  | **9 Body weight* < 5% reduced** |  |
|  | **7 Body weight* 5-10% reduced** |  |
|  | **4 Body weight* 10-20% reduced or body condition score <2** |  |
|  | **0 Body weight* > 20% reduced** |  |
| **Rapid Murine Coma and Behaviour Scale** | **Gait (0-2)**  **(none; ataxic; normal)** |  |
|  | **Balance (0-2)**  **(no body extension; extends front feet on wall; entire body lift)** |  |
|  | **Motor performance (0-2)**  **(none; 2–3 corners explored in 90 seconds; explores 4 corners in 90 seconds)** |  |
|  | **Body position (0-2)**  **(on side; hunched; full extension)** |  |
|  | **Limb strength (0-2)**  **(hypotonic, no grasp; weak pull-back[front paw grasp only]; strong pull-back [active pull away, jerk away])** |  |
|  | **Touch escape (0-2)**  **(none; unilateral; instant and bilateral [in 3 attempts])** |  |
|  | **Pinna reflex (0-2)**  **(none; unilateral; instant and bilateral [in 3 attempts])** |  |
|  | **Toe pinch (0-2)**  **(none; unilateral; instant and bilateral [in 3 attempts])** |  |
|  | **Aggression (0-2)**  **(none; bite attempt with tail bleed; bite attempt prior to tail bleed [in 5 seconds])** |  |
|  | **Grooming (0-2)**  **(ruffled, with swaths of hair out of place; dusty/piloerection; normal/clean with sheen)** |  |
|  | **RMCBS Total Score (0-20)** |  |
|  | **Total (0-30)** |  |

**28-30 = Normal.**

**24-27 = Monitor at least daily, consider supportive care if appropriate.**

**17-23 = Suffering, provide relief, observe at least twice daily regularly. Seek advice from NACWO and/or NVS.**

**12-16 = Monitor every 4 hours. Provide support as directed by NACWO and/or NVS**

**<12 = Euthanise animal**

* Assessment of Weight: Weight should be measured at least once daily, at the same time every day

For experiments lasting up to 5 days, weight loss is calculated relative to the body weight immediately prior to start of the infection (or procedure if uninfected)

For experiments lasting more than 5 days, this calculation must include adjustment for the expected weight gain that would occur in normal mice – calculated from the average weight gain of healthy control mice of the same age and gender as the experimental animals.

Body condition score is determined according to the method of Ullman-Culleré and Foltz, 1999

**References: 1. Practical use of distress scoring systems in the application of humane endpoints. M.H Lloyd & S.E. Wolfensohn**

**2. Carroll RW, Wainwright MS, Kim K-Y, Kidambi T, Go´mez ND, et al. (2010) A Rapid Murine Coma and Behavior Scale for Quantitative Assessment of Murine Cerebral Malaria. PLoS ONE 5(10): e13124. doi:10.1371/journal.pone.0013124**
