## Supplementary File 12 Antibodies used for FACS for "Comparative transcriptomic analysis reveals translationally relevant processes in mouse models of malaria"

Supplementary Table 2: Antibodies used for defining WBC proportions using FACS.

| **Antibody, (catalogue number)** | **Compensation control** |
| --- | --- |
| Alexa Fluor® 488 anti-mouse/human CD11b  Clone M1/70 (101217) Biolegend | Alexa Fluor® 488 anti-mouse CD4 Clone GK1.5 (100425) Biolegend |
| APC anti-mouse Ly-6G Clone 1A8  (127614) Biolegend | APC anti-mouse CD4 Clone GK1.5  (100411) Biolegend |
| PE anti-mouse CD19 Clone 6D5 (115508) Biolegend | PE anti-mouse CD4 Clone GK1.5 (100407) Biolegend |
| Brilliant Violet 421™ anti-mouse CD4 Clone GK1.5 (100443) Biolegend | * Brilliant Violet 421™ anti-mouse CD4 acts as its own compensation control |
| Alexa Fluor® 700 anti-mouse CD8a Clone 53-6.7  (100730) Biolegend | Alexa Fluor® 700 anti-mouse CD4 Clone GK1.5 (100429) Biolegend |
| Brilliant Violet 650™ anti-mouse CD3 Clone 17A2 (100229) Biolegend | Brilliant Violet 650™ anti-mouse CD4 Clone GK1.5 (100545) Biolegend |
